# Progressive cell fate specification in morphallactic regeneration

**DOI:** 10.1101/2024.02.08.579449

**Authors:** Clara Nuninger, Panagiotis Papasaikas, Jacqueline Ferralli, Sebastien Smallwood, Charisios D. Tsiairis

## Abstract

Through regeneration various species replace lost parts of their body. This is achieved either by growth of new structures at the amputation side (epimorphosis), as is the case of axolotl limb regeneration, or through remodeling of the remaining tissue (morphallaxis), as happens in *Hydra*. Whereas work on epimorphic regeneration support a gradual proximal to distal establishment of cell identities, morphallactic regeneration is believed to rely on initial establishment of boundary conditions that organize the re-adjustment of the pattern. Performing single cell RNA sequencing during regeneration in *Hydra*, we revealed the sequence of cells’ transdifferentiation into the missing identities. We provide evidence that morphallaxis proceeds with progressive specification of cell fates, unifying its mechanism with the one found for epimorphosis.

## INTRODUCTION

Regeneration is the process by which a fully developed organism can replace missing body parts after an injury. This can occur either by growing new structures at the site of amputation (epimorphosis) or by remodeling the remaining tissues (morphallaxis) (Fig. 1A) (Morgan 1901). In biological terms, the challenge is to recreate a complete pattern after a portion has been amputated, often described as the ‘French Flag problem’ (Fig. 1B) (Wolpert 1969). In epimorphic regeneration, exemplified by limb regeneration in axolotl, missing identities are progressively rebuilt, starting from the ones immediately next to the amputation site and moving towards the distal end (Roensch, et al. 2013). In contrast, in morphallactic regeneration it is believed that “new boundary regions are first established and new positional values are specified in relation to them” (Wolpert, et al. 2019). However, sufficient experimental evidence to support this idea are missing and it remains unclear how different the two modes of regeneration really are.

**Figure 1:**
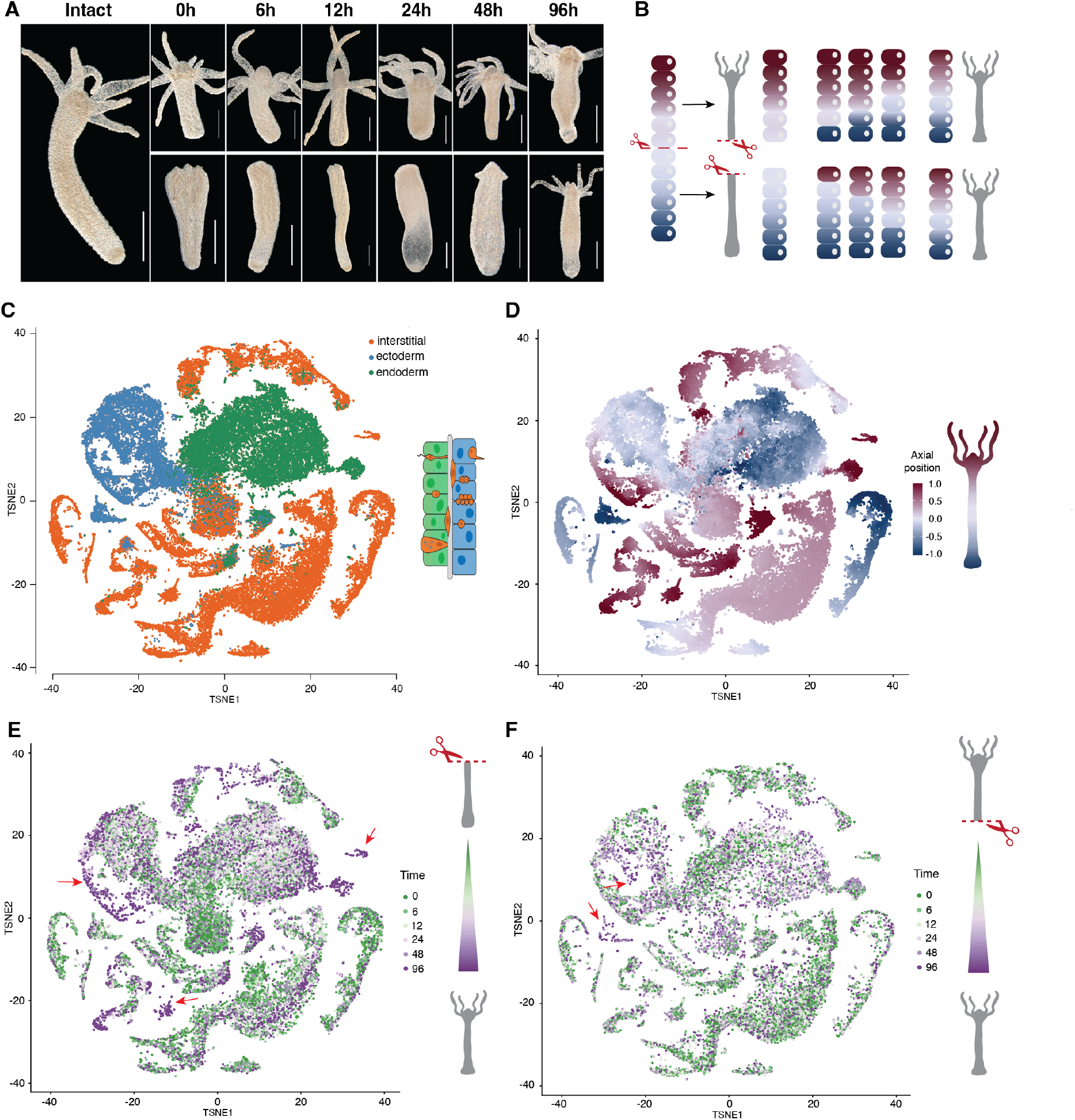
A time course scRNA sequencing of *Hydra* regeneration captures the emergence of missing axial identities. (**A**) Hydra regenerates axial pattern after mid-gastric bisection. Animals regenerating oral (lower panel) and aboral extremities (upper panel) are imaged at the indicated time points, corresponding to those used for scRNA sequencing. Scale bar = 500μm. (**B**) Schematic representation of axial differentiation during morphallaxis. Cell states along the single oral-aboral axis are color-coded from red to blue. Following bisection, boundaries are established at both amputation sites, which trigger the intercalation of missing identities. Complete patterns are restored at a smaller scale at the end of the process. (**C**) On a t-SNE projection of all sequenced cells, the main cell lineages are detected. (**D**) The axial identity of individual cell is displayed on the same t-SNE plot. Cell positions are estimated based on their gene expression profile. (**E, F**) t-SNE projections show both oral (E) and aboral (F) regeneration time courses separately. The color code indicates the temporal origin of cells. Arrows highlight cell populations with extremity identities enriched in the later time points.

*Hydra* has served as a model system for morphallactic regeneration for over two centuries (Trembley 1744). It is a freshwater polyp belonging to the phylum Cnidaria, with a tubular body organized along a single axis. Thus, a one-dimensional positional map can describe the animal pattern. At the oral extremity, tentacles that the animal uses to capture prey surround the mouth opening. At the opposite end of the axis, there is a foot structure that attaches the animal to solid substrates (Vogg, et al. 2019). The axial extremities are sources of secreted signals and are considered critical for establishing and maintaining the axial pattern (Bode 2011). The body column is composed of two epithelial layers, the outer epidermis and the inner gastrodermis. Intermingled between the epithelial cells, the specialized progeny of interstitial cells (icells) builds the nervous system, the characteristic stinging cells (nematocytes) and gametes for sexual reproduction (Holstein 2023). While the icell lineage serves essential physiological functions for the animal, it is not necessary for proper axial patterning and regeneration (Marcum and Campbell 1978).

As their mythical name implies, *Hydra* possesses remarkable regeneration abilities. When the animal is divided into pieces, each piece can regenerate into a complete, new organism. The cells that remain after amputation change their original identities to those of the missing structures without the need for increased cell proliferation or a higher cell count (Park, et al. 1970). The emergence of an oral boundary has been considered a formative step for the regeneration of the entire axial pattern (Broun, et al. 2005; Hobmayer, et al. 2000). Expression of Wnt ligands, which are constantly present at the oral end (the hypostome), has been observed at the regenerative site within a few hours after amputation (Chera, et al. 2009; Petersen, et al. 2015). It is expected that such newly formed boundaries orchestrates the scaling process of the remaining tissue. Nevertheless, the intermediate patterns that eventually bring forth the complete axial pattern are unknown.

## RESULTS

### De-novo formation of cellular identities during Hydra regeneration is revealed at a single cell resolution

To investigate the morphollactic mechanism of *Hydra*, we bisected animals and collected samples at different time points, including immediately after bisection, and at 6, 12, 24, 48, and 96 hours (Fig. 1A). Notably, for each time point, we processed the oral and aboral halves separately. These halves correspond to the regions that will regenerate a new foot or a new head structure, respectively. Following the bisection, it is assumed that terminal boundaries quickly emerge at the regenerative extremities. The proximity of these emerging boundaries to tissues that are not normally adjacent creates a discontinuity in the axial pattern. This discontinuity is hypothesized to trigger the intercalation of missing axial identities and, subsequently, the re-establishment of the complete pattern (Fig. 1B) (Bryant, et al. 1981).

Multiple animals from the corresponding time points were dissociated into single cell suspensions for subsequent single-cell RNA sequencing (sc-RNA seq). For each time point, we successfully collected approximately 4000 single-cell transcriptomes, with a median of 8000 unique molecular identifiers (UMIs) and the detection of 2240 genes per cell (Table S1). Our approach allowed us to identify cells belonging to ectodermal, endodermal, and interstitial cell lineages. Furthermore, we were able to identify all the cell types previously characterized in a single-cell transcriptome analysis of homeostatic *Hydra* (Fig. 1C and fig. S1, A and B) (Siebert, et al. 2019). During our investigation into the gene expression profiles of these cells, we specifically focused on mitotic activity to examine the origin of cells for the regeneration process. We observed cells undergoing cell division, and this phenomenon was predominantly found within the interstitial cell lineage (Fig. S1C). It is important to highlight that we did not observe significant amputation-stimulated cell division in the epithelial cell lineages, which is in line with the morphallactic regeneration process in *Hydra*.

Our research aimed to investigate the gradual restoration of the complete axial pattern during Hydra regeneration. To achieve this, we utilized a previously published positional transcriptome, which allowed us to identify gene expression profiles that vary along the oral-aboral axis (Ferenc, et al. 2021). We employed the expression of these genes as a spatial reference, essentially creating a “spatial ruler”. With this tool, we assigned each individual cell a specific positional value. This approach enabled us to establish a pseudo-axis, which is visually represented on the t-distributed stochastic neighbor embedding (t-SNE) projection. The resulting t-SNE projection revealed distinct domains with cellular identities associated with either head or foot extremities (Fig. 1D). This is consistent with the presence of the most differentiated cells with specialized functions in the terminal oral and aboral regions. Conversely, body-related cells, that are not yet committed to a fate, are more abundant and broadly distributed.

Concurrently, we were interested in understanding the temporal origin of each cell in the context of two distinct regenerative processes, specifically when either an oral or an aboral extremity is regenerating (Fig. 1, E and F). In samples undergoing head regeneration, we observed that cells with the most oral identities were detected later in the time series (Fig. 1, D and E). Conversely, in samples regenerating the foot, we found that axial identities resembling the aboral region became more prominent at later time points (Fig. 1, D and F). This temporal pattern aligns with our observation of differentiated cell types, such as specific tentacle neurons or basal disk cells of the aboral extremity, which were only present at 96 hours post-amputation (hpa) (Fig. S1, A, B and D, cluster 20 and 5). Altogether, our data allow us to follow the emergence of the missing structures at a cellular level throughout the regeneration process.

### The final boundaries are progressively established over time

To gain a deeper understanding of how cell identities change over time, we conducted separate analyses for the three cell lineages. We placed particular focus on the epithelial layers, as they are central to the primary morphallactic regeneration program (Marcum and Campbell 1978). When we projected the epidermal cells onto a t-SNE plot, we observed an arrangement of the cells according to their axial identity, with oral and aboral cells occupying diametrically opposed locations (Fig. 2A and fig. S2). Importantly, both the hypostomal and foot extremities, which are expected to play key roles in orchestrating the regeneration process, were exclusively populated by cells from the later time points (Fig. 2, B and C). This same pattern was consistent across the endodermal and interstitial lineages (Fig. S3, A and D).

**Figure 2:**
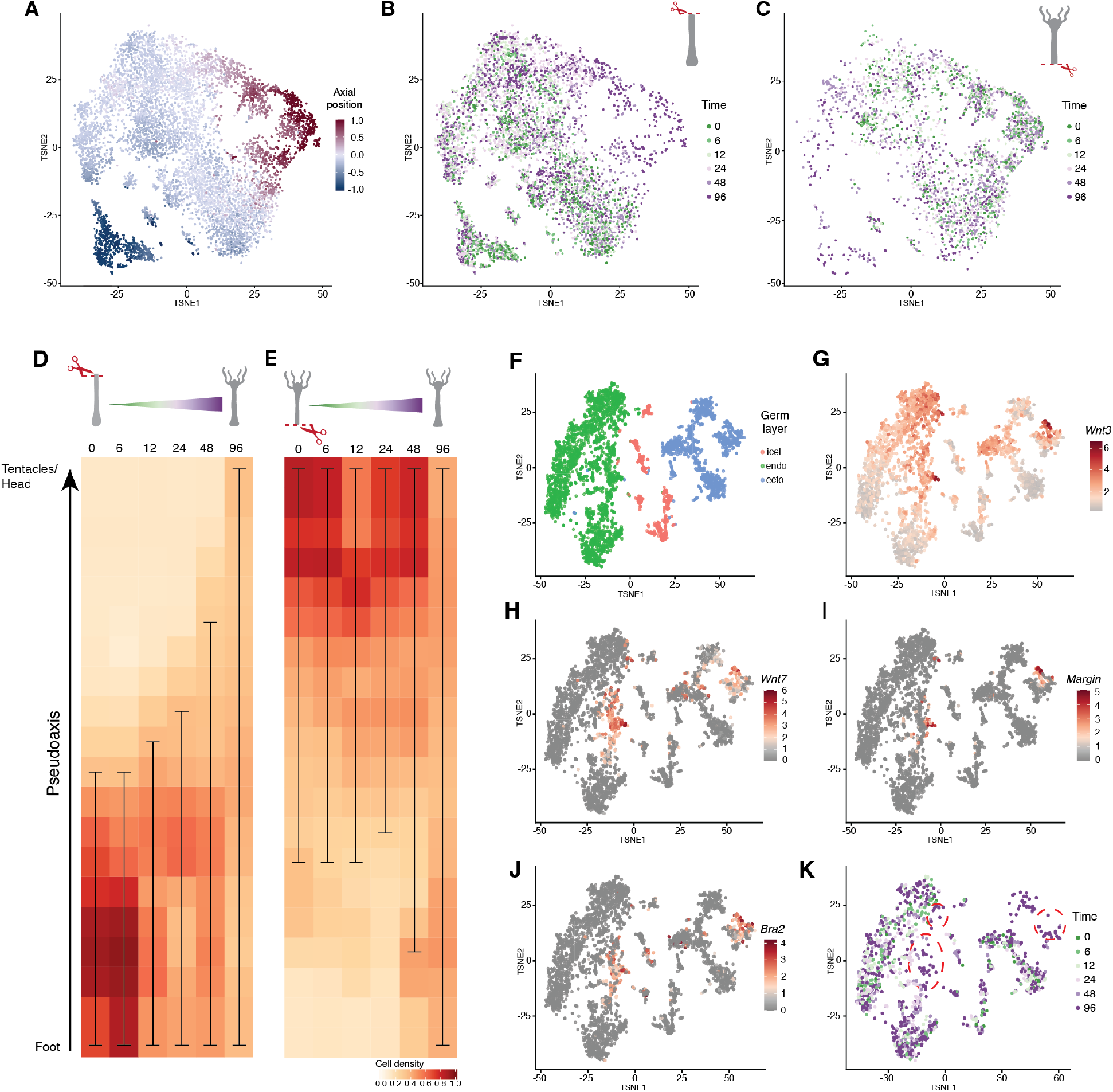
Emergence of new identities precedes the complete formation of boundaries. (**A**) A t-SNE projection displays cells from the epidermal lineage according to their oral (red) to aboral (blue) identity. (**B, C**) Cells participating in oral (B) or aboral (C) regeneration are color-coded based on their detection time. Most differentiated identities emerge predominantly at the later time points. (**D, E**) The density of epidermal cells assigned to a specific axial position in head (D) or foot (E) regenerating samples reveals a gradual acquisition of distal identities over time. The oral-aboral axis is divided into 20 bins, with segments at each time point delineating the domain encompassing more than 3% and less than 97% of the cell populations. (**F**) A t-SNE projection displays the distribution of *Wnt3* expressing cells across all cell lineages. (**G**) *Wnt3* expression levels vary, with small subpopulations exhibiting high expression in endoderm and ectoderm cells. (**H-J**) Markers associated with the hypostomal identity -*Wnt7* (H), *Margin* (I) and *Bra2* (J)-are exclusively expressed within domains of highest *Wnt3* expression, thereby defining the hypostomal boundary. (**K)** *Wnt3*^+^ cells from oral regenerating samples are plotted onto the same t-SNE dimensions and are color-coded according to their temporal origin. Hypostomal cells, enclosed in red circles, emerge at the end of the regenerating process.

To track the progression of axial differentiation, we quantified the proportion of cells associated with each axial position over time (Fig. 2, D and E). As expected, immediately following bisection, the regenerative lower fragment consisted solely of cells assigned to mid-body to foot identities, while all identities above the amputation site were absent. Nevertheless, as the regeneration progressed, the tissue gradually differentiated into new oral identities, initiating this process around 12 hpa and fully restoring the complete pattern by 96 hpa (Fig 2D). A similar, albeit slightly delayed, process occurred during aboral regeneration, with a gradual shift in cell identities toward the foot extremity occurring after 24 hpa (Fig. 2E). The sequential acquisition of new axial identities, starting with those closest to the amputation site and progressing toward more distant positions, was detected in all cell lineages (Fig. S3, G and H). This remodeling process suggests that instead of rapidly establishing new hypostomal or foot boundaries and building the body axis around them, the pattern is slowly and progressively restored toward those boundaries.

The early formation of the hypostomal boundary has long been linked to the expression of *Wnt3* at the amputation site (Hobmayer, et al. 2000; Lengfeld, et al. 2009). To better understand when this boundary forms, we conducted a thorough analysis of Wnt3-expressing cells. We found such cells in all three cell lineages, but the epithelial layers had higher expression levels(Fig. 2, F and G). Among the epithelial cells there is also considerable variation in expression levels with some populations of epidermal and gastrodermal cells showing very high expression levels. These cells co-express typical hypostomal genes including *Wnt7, Margin* and *Bra2* (Fig. 2, G to J) (Bielen, et al. 2007; Lengfeld, et al. 2009; Reddy, et al. 2019). When examining the timing of their emergence, we observed that these cells are present only at the latest time points (Fig. 2, G and K). Additionally, other genes like *Wnt16* or *Wnt9/10a*, reported at the terminal extremity, are expressed in the same high *Wnt3* domains. However, genes expressed in the oral end but not at the hypostomal boundary, such as *Alx* (Smith, et al. 2000) and *Chd* (Rentzsch, et al. 2007), are absent in these domains (Fig. S3 and Table S2). In summary, while Wnt ligand expression is detectable early during regeneration, the definitive hypostomal cells, characterized by high *Wnt3* expression and co-expression of other hypostomal genes, become detectable only towards the conclusion of the regeneration process.

### The amputation site progressively differentiates into the missing structures

Following these results, our objective was to confirm our findings by examining the spatiotemporal expression of *Wnt3* in relation to specific axial identity limits. We utilized *in situ* hybridization to concurrently observe the expression of *Wnt3* and *Nas-15*, an upper-body marker identified from a positional sequencing dataset (Ferenc, et al. 2021) (Fig. 3A and Table S2). Under normal conditions, the expression domains of the two markers along the body axis are mutually exclusive (Fig. 3B). The oral limit of *Nas-15* expression domain appears below the area of *Wnt3* expression, and the aboral limit in the middle of the gastric region.

**Figure 3:**
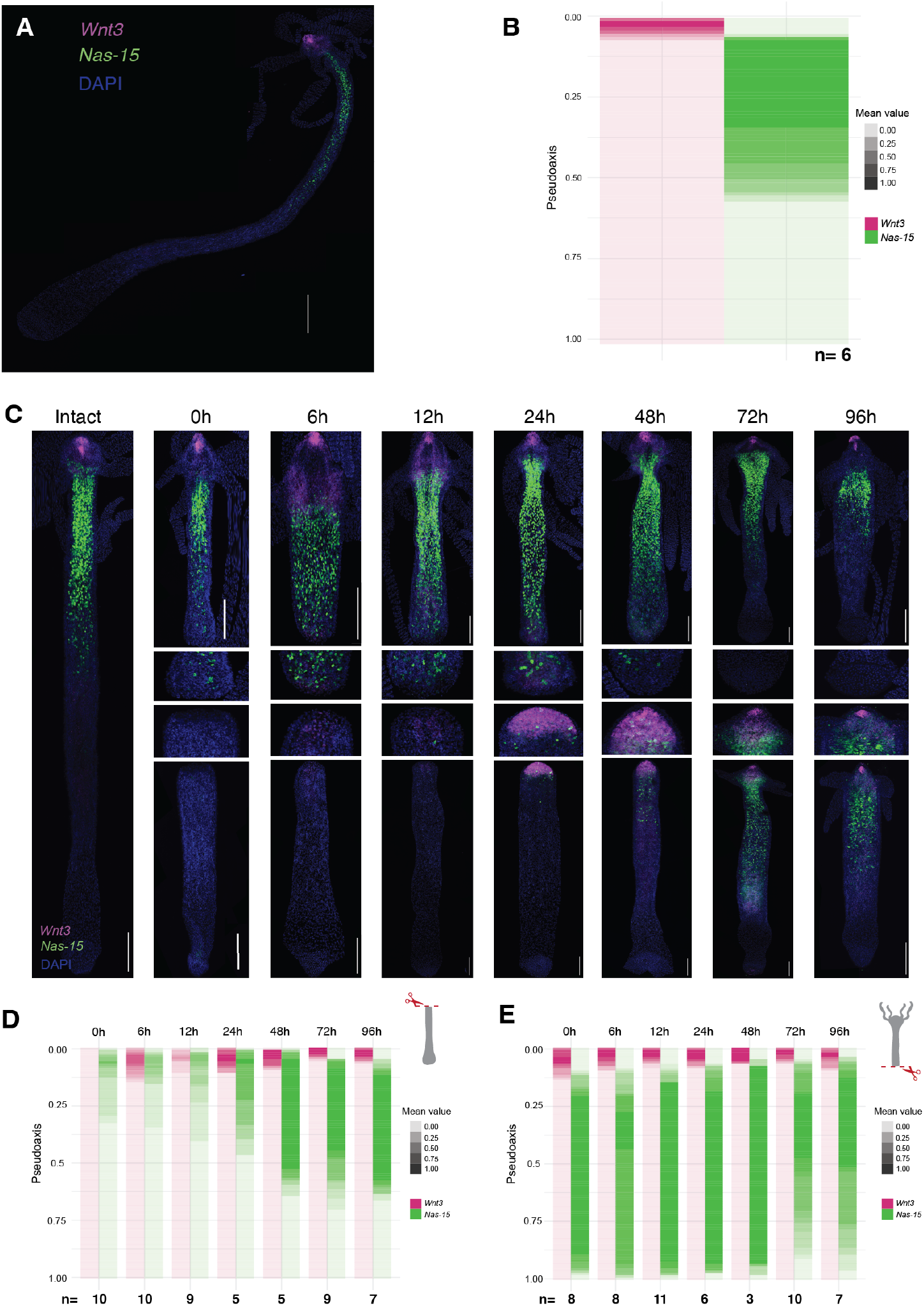
The axial pattern is progressively restored, starting from the amputation site. (**A**) In an intact *Hydra*, the expression pattern of *Wnt3* and *Nas-15* is unveiled using HCR *in situ* hybridization. Scale bar = 500μm. (**B**) The limits of *Wnt3* and *Nas-15* expression domains along the body axis indicate a distinct separation. *Wnt3* localizes at the most oral extremity, while *Nas-15* is detected in the upper body half. The average signal areas from 6 animals, normalized for axial length, are represented. (**C**) Dynamic patterns of *Wnt3* and *Nas-15* are captured at various time points during the regeneration process. Upper row shows aboral regenerating animals, lower row oral regenerating ones, and mid rows show zoom-ins on the regenerating tips. Scale bar = 200μm. (**D, E**) Quantification of *Wnt3* and *Nas-15* expression domains during oral (D) and aboral (E) regeneration illustrates the gradual changes of axial patterns over time, progressing from the amputation site. The sample numbers (n), used to average the expression areas along the normalized axis, are indicated for each time point.

During the regeneration of the oral extremity, we noted the early expression of *Wnt3* at the amputation site, with no *Nas-15* signal detected (Fig. 3, C and D). This early Wnt signaling, also observed in aboral regeneration, has been previously reported as part of the rapid wound response following damage (Cazet, et al. 2021; Chera, et al. 2009; Gufler, et al. 2018; Tursch, et al. 2022). *Nas-15*^+^ cells emerge in the oral region starting from 12 hpa and progressively expand towards the aboral end as regeneration progresses. Intriguingly, after 72 hours of regeneration, *Wnt3* expression becomes localized, resembling a hypostomal boundary, while the *Nas-15* domain is gradually excluded from the oral end. Thus, although *Wnt3* is expressed early, the surrounding cells progressively acquire their terminal identities Similarly, during the regeneration of the aboral part, there is an increasing distance of the *Nas-15* domain from the regenerating end, while the *Wnt3* area shrinks to accommodate the smaller but complete animal pattern at the end of regeneration (Fig. 3, C and E). These findings suggest that the regenerative oral and aboral regions are initially not domains with fixed identities but rather represent dynamically differentiating endpoint structures.

To investigate whether tissue differentiation initiates in the regenerative extremities without immediate boundary formation, we conducted a detailed examination of the tissue identity beneath the amputation site. The regenerating tip at the oral end will definitely undergo identity change during regeneration as it will transform into a new hypostome surrounded by tentacles. We systematically collected and RNA-sequenced samples from the oral end of regenerating animals at one-hour intervals over a 48-hour period. The samples were then ordered based on pseudo-time to account for their asynchronous regeneration rates (Fig. 4A), allowing us to monitor the dynamic expression profile of the regeneration front. At the same time, we utilized a previously acquired positional transcriptome (Ferenc, et al. 2021) to generate a principal component analysis (PCA) projection map of axial identities (Fig. 4B). The projection of pseudo-time-ordered regenerative tips onto the same PCA dimension revealed that initially, the samples clustered alongside mid-body fragments, as expected following bisection (Fig. 4C). As regeneration progressed, we observed a gradual transition of the amputation site from mid-body towards a more head-like identity, passing through intermediate axial states. By 48 hours, the amputation tips closely approached head fragments without fully reaching them, aligning with our previous observations (Fig. 2D and Fig. 3, C and D). These results make evident that regeneration does not commence with the immediate establishment of a boundary.

**Figure 4:**
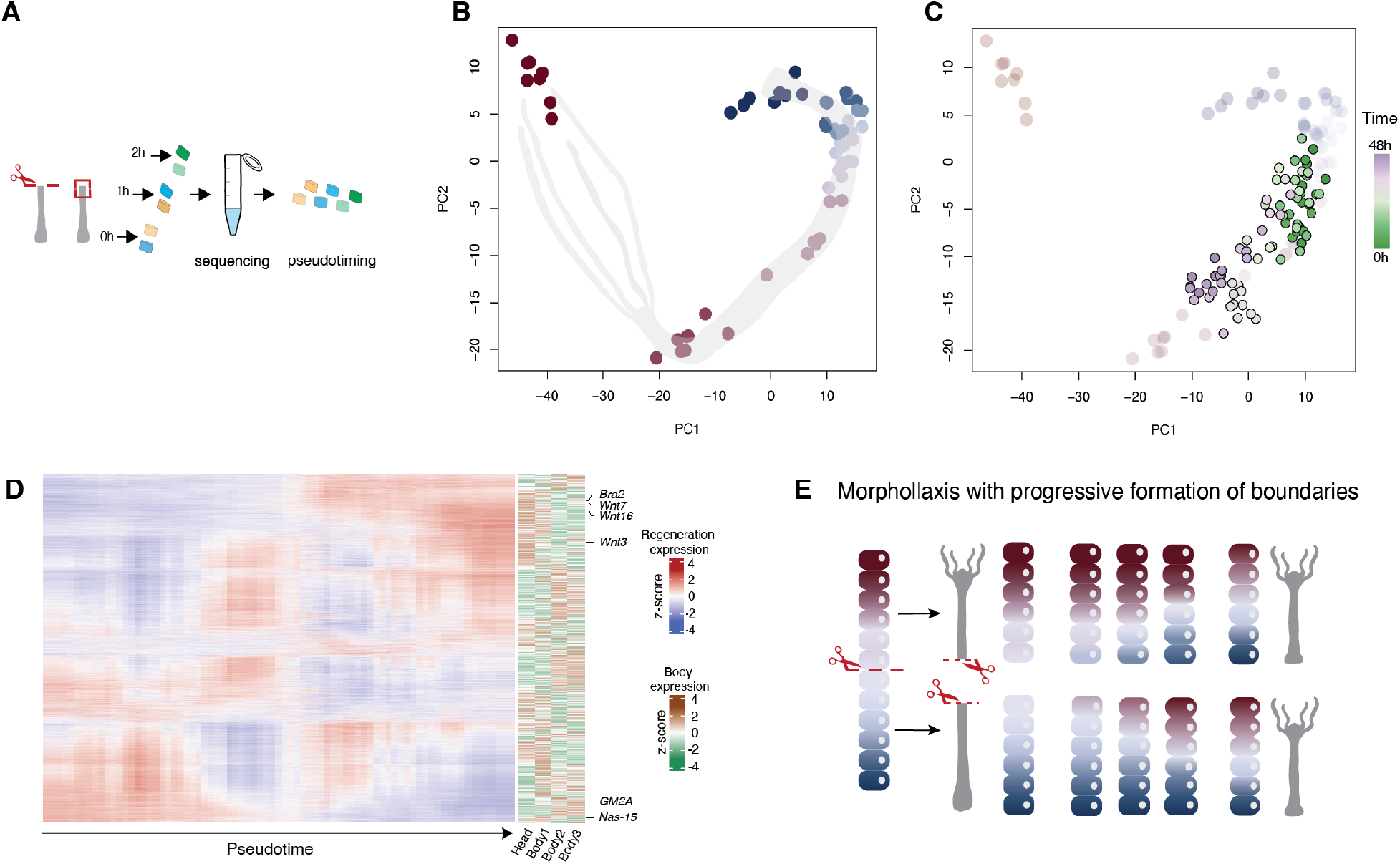
The development of distal structures in the regenerative tip initiates from the nearest missing axial neighbor. **(A**) Schematic of the experimental design. Regenerating tips from two different animals per time point, are collected at hourly intervals over a 48-hour time course, processed individually for RNA sequencing and pseudo-time ordered. (**B**) Body segments acquired along the oral-aboral axis were sequenced (in Ferenc et al., 2021) and projected onto a principal components analysis (PCA) space, revealing a map of positional identity. The color-coding indicates the axial origin of the body segments, transitioning from tentacles (red) to foot (blue). Eight replicates per body segment are represented. (**C**) Projection of the pseudo-time ordered regenerative tips onto the PCA space, defined by the expression patterns of body segments, unveils a gradual shift, from early (green) to late (purple) time points, of the tip’s identity towards the oral end. (**D**) Heatmaps of differentially expressed genes over the regeneration time course (left) and their respective spatial distribution (right). Z-scaled log expression values are presented. Highlighted are examples of genes that are expressed early and then switch off (e.g., *GM2A*), or emerge later on (e.g., *Wnt16*). (**E**) Morphallactic regeneration happens with a progressive acquisition of identities from the nearest missing neighbor towards the most distal ones. The end of the process is marked by the delineation of axial boundaries. Cell states along the single oral-aboral axis are color-coded from red to blue.

The hypostomal boundary represents a confined domain of the *Hydra* body, with a distinct gene expression profile. To trace the temporal sequence of its formation, we scrutinized the differentially expressed genes over time, emphasizing two specific patterns: some genes are initially expressed at the regenerative tip and later downregulated, while others are only expressed later in the time course (Fig. 4D). Notably, *Wnt3* follows a periodic expression pattern, peaking for a first time shortly before the midpoint of the time course. In contrast, many definitive markers of the terminal end, such as *Wnt7* and *Bra2*, are expressed later, aligning with our earlier findings that the complete hypostomal boundary is established towards the end of regeneration (Fig. 2, G to K). Conversely, genes characteristic of the middle and upper-half of the gastric region, such as *Nas-15* and *GM2A*, are upregulated early, with their expression decreasing towards the end of the time course (Fig 4D and Table S2). Taken together, these findings suggest that the hypostomal boundary emerges gradually over time rather than being specified early on to guide the formation of other missing structures (Fig. 4E).

## DISCUSSION

Our primary discovery contradicts widely held theories about the mechanism underlying morphallactic regeneration. Previously, it was believed that the early establishment of boundary identities at the amputation site would create a disruption in the sequence of cell fates, leading to the insertion of missing positions between adjacent points (Bryant, et al. 1981). However, our data reveal that in *Hydra*, cells gradually establish missing positional identities, beginning with those closest to the amputation site, and the regeneration process is completed when the boundary is reached. This sequential establishment of cell identities, without the emergence of identity gaps, mirrors what is observed in epimorphic regeneration (Roensch, et al. 2013). Despite differences in the requirement for cell proliferation, both epimorphosis and morphallaxis adhere to the same underlying principle.

Our research provides a framework to help us understand previous observations about the varying speed of regeneration completion. Previous studies have shown that head regeneration is faster when the amputation site is closer to the oral boundary (Mookerjee and Bhattacharjee 1967). This happens because the regenerating tip has a shorter distance to traverse through a smaller set of positional identities to ultimately reach the oral extremity. Furthermore, researchers have noticed the temporary expression of several genes, including *Alx, Zic4, Ks1*, and *Chd*, at the regenerating tip (Endl, et al. 1999; Rentzsch, et al. 2007; Smith, et al. 2000; Vogg, et al. 2022). However, these genes are typically expressed below the oral tip of the animal. We can interpret this expression as the regenerating tip acquires these identities before ultimately adopting the hypostomal, or oral end, identity. Nevertheless, a clear marker for the hypostomal identity, *Wnt3*, is expressed very early at the head amputation site (Lengfeld, et al. 2009). Our data reveals differences not only in the levels of *Wnt3* expression over time but also in the set of genes co-expressed with *Wnt3*. Genes that clearly mark the definitive hypostomal end are co-expressed at a later stage in the regeneration process. The continuous presence of Wnt signaling, a known requirement for regeneration (Broun, et al. 2005), appears to fuel the differentiation process and might even be a temporarily integrated signal for the cells at the regenerating tip (Briscoe and Small 2015), as they transition in their positional identity over time.

The mechanism we have presented, along with the associated data, can serve as a valuable guide for future research into pattern formation. In addition to complementing recent studies that have compared head and foot regeneration (Cazet, et al. 2021), our work provides both a resource and a framework for investigating wound-triggered differentiation and the effects of morphogen signals at the cellular level. Morphallaxis essentially involves a scaling process, and adjustments in morphogen gradients have been linked to scaling phenomena (Čapek and Müller 2019). As we have demonstrated here, the regeneration process in *Hydra* provides a reliable system to test these various paradigms.

## MATERIAL AND METHODS

### Animal strains and culture conditions

All experiments were performed with strains of *Hydra magnipapillata*. The cultures were maintained in Hydra medium (HM) [1 mM CaCl_2_, 0.2 mM NaHCO_3_, 0.02 mM KCl, 0.02 mM MgCl_2_, and 0.2 mM tris-HCl (pH 7.4)] at 18°C and fed three times per week with freshly hatched *Artemia nauplii*. For the experiments, individuals without buds and starved for at least 24 hours were used. All experimental procedures adhered to ethical standards and complied with Swiss national regulations for the use of animals in research.

### Hybridization chain reaction RNA-FISH

*In situ* hybridization was performed according to previously published protocol (Vogg, et al. 2022) with minor modifications. Briefly, *Hydras* were relaxed in 2% urethan/HM for 1min before fixation for 30min to 1h in 4% paraformaldehyde (PFA)/PBS (v/v). The fixative was washed away with two washes of 10min in PBS/0.1% Tween 20 (PBST). Tissues were rehydrated with an incubation of 1h in a 70% ethanol/PBST (v/v) solution, followed by 5min in a 35% ethanol/PBST (v/v) solution at 4°C. Samples were washed twice 10min with PBST before incubation in 250μl of HCR probe hybridization buffer, without probe, for 30min at 37°C. Samples were then incubated with 250μl of HCR probe hybridization buffer containing 1.5pmol of *Wnt3* and *Nas-15* hybridization probes overnight (12 to 16 hours) at 37°C. The probes were synthesized by Molecular Instruments (lot number PRK123, PRK125). Hybridization probes were washed out with 5 washes of 1h in 500μl of HCR probe wash buffer at 37°C, followed by two washes of 5min with a 5× sodium chloride sodium citrate/0.1% Tween 20 (SSCT) solution. Signal was pre-amplified with 250μl of amplification buffer for 30min. 7.5pmol of amplifying hairpin RNAs, provided by Molecular Instruments, were heated at 95°C for 1min 30s and allowed to cool down in the dark for 30min. The preamplification solution was replaced by 250μl of amplification buffer containing the two hairpins, and samples were incubated overnight (12 to 16 hours) in the dark. Excess of probe was washed out with several washes in 5× SSCT solution in the dark: twice 5min, three times 30min, once 30min with 4′,6-diamidino-2-phenylindole (DAPI, 1μg/ml), and once 5min. Samples were mounted on a slide with ProLong Gold Antifade™ for imaging.

### Preparation of single-cell suspension of regenerating *Hydras* for RNA sequencing

20-40 *hydras* were cut in half per time point. Both upper and lower parts were allowed to regenerate in HM before being separately collected in 1.5mL tubes. Animals were starved 4 days prior dissociation. Samples were incubated in 1mL of Protease solution composed of 7.5mg/mL of protease E (P8811, Sigma Aldrich) diluted in dissociation medium [6mM CaCI_2_, 1.2mM MgSO_4_, 3.6mM KCl, 12.5mM *N*-tris[hydroxymethyl]methyl-2-aminoethanesulfonic acid (TES), 6mM sodium pyruvate, 6mM sodium citrate, 4mM glucose (pH 6.9)] for 1h at 24°C, 300rpm on a thermoblock, protected from light. Animals were dissociated into single cells mechanically by pipetting 20 times up and down using a 1000μL pipette. Cells were strained through a sterile CellTrics® 30μm cell strainer (04-004-2326, Sysmex), collected in a centrifuge tube and spin 5min, 300xg at 4°C. Supernatant was removed and cells were resuspended in 1mL of dissociation medium + 0.01% Bovin serum albumin (BSA) (A3912, Sigma Aldrich). Cells were centrifuged again 5min at 300xg 4°C and resuspended in a 1mL dissociation medium + 0.01% BSA. Cells were strained through a 30μm cell strainer and collected in a FACS tube. Dead cells were stained by adding 5μL of DRAQ7™ (D15106, Thermofisher) to the suspension.

### Fluorescence activated cell sorting (FACS)

Dissociated cells were sorted on a BD FACS Aria cell sorter with a 70μm nozzle. Cells were sorted based on the DRAQ7™ staining (Alexa fluor 700) and according to the cell diameter based on the forward scatter (FSC). 8000 cells of all cell types and 8000 of enriched epithelial cells were sorted and collected in the same collection tube in a 0.04% BSA/ PBS (without CaCl_2_ and MgCl_2_) solution for all time points. Post-FACS survival assays, after incubation on ice for 30min, were performed to ensure that *Hydra* cells were still viable.

### 10x scRNA sequencing

FACS-sorted cells (16K) were immediately processed for scRNA-seq using 10X genomics Chromium Single Cell 3’ Reagents v3.1 with UDI indexes, according to the manufacturer’s protocol. Final libraries were sequenced on an Illumina NovaSeq6000 platform, 2x56 paired-end cycles. Fastq files were generated using Illumina bcl2fastq2 pipeline and further processed with 10X genomics cellranger.

### RNA-seq time course of regenerating tip extremity

Over a period of 12 hours, multiple animals per hour were cut in half and let regenerate in HM. After the last bisection, the oral ends of two regenerating *Hydras* were collected per time points, creating a 12-hour time course. This sequence was replicated to extend the observation period to 48 hours. Each fragment was directly lysed in 90μL of RLT buffer (Single Cell RNA purification kit, Norgen Biotek) containing 1% of β-mercaptoethanol. Subsequently, the lysed samples were rapidly frozen on dry ice and stored at −80°C for later RNA isolation. RNA was purified following the manufacturer’s protocol. RNA quality was assessed using an Agilent Bioanalyzer and approximately 5ng was used as input to generate RNA sequencing libraries using SmartSeq2, as previously reported (*17*). While all samples were processed in parallel, time points 0 to 23 hours were first sequenced on a HiSeq2500, 50 cycles single-ended; Time points 24h to 47h were sequenced at a later time on a NovaSeq6000, 2x56 paired-end cycles, after treatment of the library pool with Illumina free adapter blocking reagent to reduce rates of index hoping. The change of sequencers was the result of the management of the life cycle of the instruments by the FMI genomics platform.

### Layer classification, axial and time ordering and visualization of RNAseq samples

Quantification of the generated single cell libraries was performed using the Salmon-Alevin software suite (Salmon version 1.6.0) against the ncbi Hydra 102 transcriptome. Example command:

*salmon alevin -l ISR -1 $R1 -2 $R2 --chromiumV3 -i $reference -p $threads -o OUTDIR --tgMap*

*$tx2genemap --dumpFeatures --numCellBootstraps 50 --expectCells 10000*

QC metrics for cells were generated using the scater addPerCellQC function. Cells with < 750 detected genes were removed from subsequent analyses. After removal of low quality cells we recovered a total of 53109 cells corresponding to approximately 4000 cells per time point after amputation for each regeneration experiment (see Table S1).

### Data integration of generated libraries and visualization

Single cell data from four different libraries were integrated in the complete single cell dataset. Scaling normalization within each batch was performed using the multiBatchNorm function from the Bioconductor batchelor package.

For selection of variable genes, expression variance was model using the modelGeneVar function from the scan package separately for each batch and subsequently the variance decompositions were combined using the scran combineVar function. Variable genes were defined based on the obtained variance modelling statistics using the scran getTopHVGs function (parameter prop=0.75).

Combined projections of batches and germ-layer specific projections were obtained using the fast mutual nearest neighbors method (fastMNN) from the Bioconductor batchelor package. Projections reflecting axial distribution of cells in Hydra were obtained using as input to dimensionality reduction the expression matrix of the intersection of variable and axial graded genes (see following section).

In all cases default sets of parameters were used unless otherwise indicated.

### Cell germ-layer classification, pseudo-axial cell identities, cell mitotic state assignment, and body segment markers

Cell germ layer classification uses a simple three-way classifier that assigns class according to the max of the mean normalized, scaled summed counts of interstitial, epidermal and endodermal markers from (Siebert, et al. 2019).

Pseudo-body axial identity was derived, independently for each layer, using a collection of axial-graded genes that show strong positive or negative association (|ρ| > 0.7) with the oral-aboral axis according to (17). A single axial index was obtained using the difference of the mean log transformed expression values for the positive and negative axial gene sets. The numeric index was finally rescaled in the [-1, 1] range.

Mitotic cell states were assigned using a set of the mouse mitotic cell markers from the MSigDB database (GOBP_MITOTIC_CELL_CYCLE collection) that were mapped to orthologous Hydra genes according to ncbi annotation. Mitotic indices were obtained using the mean log-transformed expression values of the mitotic cell markers.

Both the derived axial indices and mitotic state indices of cells were smoothed using the averages across k-nearest-neighbors using a narrow kernel (k=10).

Individual body segment markers were identified from body segment expression data taken from(Ferenc, et al. 2021). Candidate markers for Tentacles, Hypostome, Upper and Lower Body and Foot were identified as genes with maximal expression value in the corresponding segment. Subsequently germ-layer specific markers for each segment were defined as the subset of the candidate marker genes for the corresponding segment that show maximal expression in cells with axial identities consistent with that segment. In turn, axial patterns of each candidate gene and for each germ layer were generated using the average gene expression in 20 axial bins.

### Pseudo-time ordering and projection of the bulk RNA-seq samples for tip regeneration

Quantification of the bulk RNAseq libraries for tip regeneration was performed using the STAR software (version 2.5.0) and the NCBI Hydra 102 transcriptome for gene count generation.

Pseudo-time ordering of the bulk RNA-seq samples from the tip regeneration experiment was performed using a 1D projection on the unique principal component that is significantly associated (r^2^ = 0.34, p-value < 1e-16) with the time-of-collection after amputation value. PCA projection on a space that reflects body segment identity was performed as described in (Ferenc, et al. 2021).

### Image acquisition

Brightfield images were acquired with a stereo microscope composed of a Leica DFC 480 camera and a Plan APO 1.0x objective (Leica 10450028). Images were acquired with the IMS software.

Fluorescent imaging was conducted using a spinning disc confocal microscope comprising a Yokogawa CSU W1 scanning unit with Dual T2, coupled with an inverted motorized stand Zeiss AxioObserver7. The system equipped with a back-illuminated Prime 95B sCMOS camera (1200 pixels × 1200 pixels, Photometrics) and is configured with four excitation lasers operating at 405nm, 488nm, 561nm and 640nm. The imaging objective employed was a Plan Apochromat 20×/0.8 WD 0.55 (Zeiss 420650-9901). Image acquisition was performed using Visiview software from Visitron.

Multiples images were taken for each animal, covering both the xy and z dimensions. The z-dimension images were obtained with slices of 1μm thickness. Regenerating feet and heads of the same experimental day were acquired with the same exposure time and laser intensity, adjusted according to the corresponding regenerating foot condition where both markers are expressed from the beginning of the time course.

### Quantification of fluorescent signal domains

Images were processed and analyzed on ImageJ/Fiji (v. 2.1) and minimal-maximum intensities were adjusted according to the foot regenerating condition, characterized by constitutive expression of both axial markers. This adjustement approach was applied uniformly to samples from the same experimental day. For each animal, images along the XYZ dimensions were stitched together, using the corresponding imageJ plugin. Additionally, a maximum projection of the Z-stacks was perfomed using the built-in Z-project function. A manually drawn segmented line, measuring 1500μm in thickness, extended from the oral to the aboral extremity of the animal. Subsequently, the image was straightened using the straigten function along the delineated line. The signal area boundaries for *Wnt3* and *Nas-15* were manually defined along the pseudoaxis and the channels were then merged for visualization. Broken or folded animals were withdrawn from the analysis.

## Acknowledgments

We are grateful to the FMI Imaging, Genomics and Cell Sorting facilities for their support with experiments. The analysis of the transcriptomics data was done with the support of the FMI Bioinformatics facility. We would also like to thank Helge Grosshans, Elly Tanaka, and all members of our lab for helpful comments on the manuscript.

## Notes

### Competing Interest Statement

The authors have declared no competing interest.

## REFERENCES

Bielen, H., et al. 2007 Divergent functions of two ancient Hydra Brachyury paralogues suggest specific roles for their C-terminal domains in tissue fate induction. Development 134(23):4187–97.

Bode, Hans 2011 Axis Formation in Hydra. Annual Review of Genetics 45(1):105–117.

Briscoe, J., and S. Small 2015 Morphogen rules: design principles of gradient-mediated embryo patterning. Development 142(23):3996–4009.

Broun, M., et al. 2005 Formation of the head organizer in hydra involves the canonical Wnt pathway. Development 132(12):2907–16.

Bryant, S. V., V. French, and P. J. Bryant 1981 Distal regeneration and symmetry. Science 212(4498):993–1002.

Čapek, D., and P. Müller 2019 Positional information and tissue scaling during development and regeneration. Development 146(24).

Cazet, J. F., A. Cho, and C. E. Juliano 2021 Generic injuries are sufficient to induce ectopic Wnt organizers in Hydra. Elife 10.

Chera, S., et al. 2009 Apoptotic cells provide an unexpected source of Wnt3 signaling to drive hydra head regeneration. Dev Cell 17(2):279–89.

Endl, I., J. U. Lohmann, and T. C. Bosch 1999 Head-specific gene expression in Hydra: complexity of DNA-protein interactions at the promoter of ks1 is inversely correlated to the head activation potential. Proc Natl Acad Sci U S A 96(4):1445–50.

Ferenc, J., et al. 2021 Mechanical oscillations orchestrate axial patterning through Wnt activation in Hydra. Sci Adv 7(50):eabj6897.

Gufler, S., et al. 2018 β-Catenin acts in a position-independent regeneration response in the simple eumetazoan Hydra. Dev Biol 433(2):310–323.

Hobmayer, B., et al. 2000 WNT signalling molecules act in axis formation in the diploblastic metazoan Hydra. Nature 407(6801):186–9.

Holstein, T. W. 2023 The Hydra stem cell system - Revisited. Cells Dev 174:203846.

Lengfeld, T., et al. 2009 Multiple Wnts are involved in Hydra organizer formation and regeneration. Dev Biol 330(1):186–99.

Marcum, B. A., and R. D. Campbell 1978 Development of Hydra lacking nerve and interstitial cells. J Cell Sci 29:17–33.

Mookerjee, S., and A. Bhattacharjee 1967 Regeneration time at the different levels of hydra. Wilhelm Roux Arch Entwickl Mech Org 158(3):301–314.

Morgan, T.H. 1901 Regeneration: Macmillan.

Park, H. D., A. B. Ortmeyer, and D. P. Blankenbaker 1970 Cell division during regeneration in Hydra. Nature 227(5258):617–9.

Petersen, H. O., et al. 2015 A Comprehensive Transcriptomic and Proteomic Analysis of Hydra Head Regeneration. Mol Biol Evol 32(8):1928–47.

Reddy, P. C., et al. 2019 Molecular signature of an ancient organizer regulated by Wnt/β-catenin signalling during primary body axis patterning in Hydra. Commun Biol 2:434.

Rentzsch, F., et al. 2007 An ancient chordin-like gene in organizer formation of Hydra. Proc Natl Acad Sci U S A 104(9):3249–54.

Roensch, K., et al. 2013 Progressive specification rather than intercalation of segments during limb regeneration. Science 342(6164):1375–9.

Siebert, S., et al. 2019 Stem cell differentiation trajectories in Hydra resolved at single-cell resolution. Science 365(6451).

Smith, K. M., L. Gee, and H. R. Bode 2000 HyAlx, an aristaless-related gene, is involved in tentacle formation in hydra. Development 127(22):4743–52.

Trembley, A. 1744 Mémoires pour servir à l’histoire du genre de polypes d’eau douce, à bras en forme de cornes. Par M. Tremblay, de la Société Royale de Londress, etc. Tome premier [-Tome second].

Tursch, A., et al. 2022 Injury-induced MAPK activation triggers body axis formation in Hydra by default Wnt signaling. Proc Natl Acad Sci U S A 119(35):e2204122119.

Vogg, M. C., et al. 2022 The transcription factor Zic4 promotes tentacle formation and prevents epithelial transdifferentiation in Hydra. Sci Adv 8(51):eabo0694.

Vogg, M. C., B. Galliot, and C. D. Tsiairis 2019 Model systems for regeneration: Hydra. Development 146(21).

Wolpert, L. 1969 Positional information and the spatial pattern of cellular differentiation. J Theor Biol 25(1):1–47.

Wolpert, L., C. Tickle, and A.M. Arias 2019 Principles of Development: Oxford University Press.

